# Species-specific strategies increase unpredictability of escape flight in eared moths

**DOI:** 10.1101/485698

**Authors:** Theresa Hügel, Holger R. Goerlitz

## Abstract

1. Many prey species overlap in time and space and are hunted by the same predators. A common anti-predator behaviour are evasive manoeuvres to escape an attacking predator. The escape-tactic diversity hypothesis postulates that species-specific differences in evasive behaviour will increase the overall unpredictability experienced by predators within a predator-prey community. Evolutionary, escape-tactic diversity would be driven by the enhanced predator protection for each prey individual in the community. However, escape-tactic diversity could also be a functional consequence of morphological differences that correlate with evasive capabilities.
2. Echolocating bats and eared moths are a textbook example of predator-prey interactions. Moths exhibit evasive flight with diverse tactics; however, the variability of their evasive flight within and between species has never been quantified systematically. In addition, moth species show variation in size, which correlates with their flight capability.
3. We recorded flight strength during tethered flight of eight sympatric moth species in response to the same level of simulated bat predation. Our method allowed us to record kinematic parameters that are correlated with evasive flight in a controlled way to investigate species-specific differences in escape tactics.
4. We show species-specific and size-independent differences in both overall flight strength and change of flight strength over time, confirming the escape-tactic diversity hypothesis for eared moths. Additionally, we show strong inter-individual differences in evasive flight within some species. This diversity in escape tactic between eared moths increases the overall unpredictability of evasive flight experienced by bat predators, likely providing increased protection against predatory bats for the single individual.

## INTRODUCTION

To successfully escape from a predator in a chase, prey animals have two main options: being faster or being more manoeuvrable (Howland, 1974). Higher manoeuvrability allows prey to abruptly change its movement trajectory, making its behaviour variable and unpredictable. Unpredictable, erratic, or “protean” behaviour is a common escape strategy found in numerous prey taxa (Humphries & Driver, 1967, 1970). In addition to variability within an individual and a species, interspecific variability in escape behaviour has the potential to add another level of unpredictability. If multiple species in a prey community vary in the parameters of their evasive movement, the overall variability and unpredictability would increase and afford even higher protection against predators for the single individual (Schall & Pianka, 1985). Previous studies of prey communities have shown that different species can use very different anti-predator strategies (Hölker et al., 2007; Randall, Hatch, & Hekkala, 1995; Wohlfahrt, Mikolajewski, Joop, & Suhling, 2006; Wolf, 1985), such as changing microhabitat versus reducing activity in response to the same predation risk (Hölker et al., 2007). In addition, prey could also exhibit inter-specific differences within one specific anti-predator strategy, such as evasive movement (‘escape-tactic diversity hypothesis’, (Schall & Pianka, 1985)). Interspecific differences in evasive behaviour might be explained by interspecific differences in anatomy, such as muscle volume, weight and size, which are correlated with speed, acceleration and turning performance (Wilson et al., 2018).

Echolocating bats and eared moths are an ideal study system to address this question. Both groups interact in an evolutionary predator-prey arms race (Jens Rydell, Jones, & Waters, 1995; Ter Hofstede & Ratcliffe, 2016; Waters, 2003). Many insectivorous bats have a broad diet consisting of many different species of moths and other nocturnal insects (Anthony & Kunz, 1977; Bogdanowicz, Fenton, & Daleszczyk, 1999; Findley & Black, 1983), which they hunt by echolocation in mid-air (Denzinger & Schnitzler, 2013; Fenton, Portfors, Rautenbach, & Waterman, 1998). Many flying moths rely on evasive flight to escape echolocating bats. Moth evasive flight involves a two-staged response towards an attacking bat [8]. For distant bats, moths receive a faint echolocation call and steer away from the bat to avoid detection. For a close-by bat, moths receive a loud call and elicit last-ditch evasive flight to escape the bat. The last-ditch evasive flight performed by many eared moths when trying to escape an echolocating bat includes zig-zagging, loops, tight turns, passive dives, and power dives (Corcoran & Conner, 2012; Kenneth D. Roeder, 1962). Despite decades of research, this evasive behaviour was never systematically quantified and compared on a species level. Several studies observed a “general response” without going into further descriptions or quantifications of the actual behaviour (Göpfert & Wasserthal, 1999; J. Rydell, Roininen, & Philip, 2000; Svensson, Rydell, & Brown, 1999), or quantified only the consequences of anti-predator behaviour (Fullard, Muma, & Dawson, 2003; Nakano, Ihara, Mishiro, Toyama, & Toda, 2015), or lacked species identification or standardized conditions (Agee, 1969; K D Roeder, 1966; J. Rydell, Skals, Surlykke, & Svensson, 1997; Treat, 1955). Hence, it is unknown if variation in evasive flight occurs within a single individual, between individuals from the same species, or between different species. As many different moths occur in the same habitat (Conrad, Woiwod, Parsons, Fox, & Warren, 2004; Scalercio, Infusino, & Woiwod, 2008; Summerville & Crist, 2004; Truxa & Fiedler, 2012), any given bat will encounter many different moth species while hunting. Interspecific variation in evasive flight would increase the variation experienced by each bat and should provide increased protection against predatory bats for each single moth individual.

Here, we systematically quantified vertical flight strength of eight species of sympatric eared moths with different sizes during tethered flight in a flight recorder. We studied size as one explanatory variable underlying potential species-specific differences in evasive flight, since size is positively correlated with acceleration in a butterfly (Berwaerts, Van Dyck, & Aerts, 2002), and negatively correlated with manoeuvrability in insects (Dudley, 2002). We use vertical flight strength as proxy for flight speed, which, like many other variables, increases when moths actively perform last-ditch evasive flight (Corcoran & Conner, 2016, 2017). Even though tethered flight does not allow us to study actual 3D-flight behaviour, measurements of flight strength and its temporal variation do reveal species-specific strategies for and interspecific differences in evasive flight. Furthermore, tethered flight allowed us to exclude variation in received sensory input, by exposing all individuals to the same acoustic stimulus mimicking an attacking bat, to trigger last-ditch evasive flight. We quantified the inter-individual and inter-specific variability within a single anti-predator-strategy, (I) testing the hypothesis of escape-tactic diversity in moths, predicting that last-ditch evasive flight varies more between species than within species of a sympatric moth community, and (II) predicting that moth size is one explaining variable for this variation.

## METHODS

### Flight recorder and experimental setup

We developed a flight recorder to quantify moths’ vertical flight strength (Fig. 1a). The transducer consisted of two small broadband loudspeakers (25 mm nominal diameter; NSW1-205-8A, AuraSound, Guangzhou, China), which were connected via a light wooden connector that was glued onto the loudspeakers’ membranes. A plastic cylinder was centrally fixed to the wooden connector and served as a holder for an insect pin attached to a moth (see below). Vertical forces generated by the moth’s flight were transferred via the pin and the wooden connector to the membranes of both speakers, generating voltage fluctuations that were amplified and recorded via a soundcard (192 kHz sampling rate, 16 bit resolution; Fireface UC, RME, Haimhausen, Germany). The flight recorder was mounted centrally in an anechoic chamber (Desone Modular Akustik, Berlin, Germany, interior volume: 0.96×0.96×0.77m^3^, Fig 1b). A loudspeaker (NeoCD1.0 Ribbon Tweeter, Fountek Electronics, Jiaxing, China) was mounted 34 cm behind the moth and driven by a power amplifier (TA-FE330R, Sony, Tokio, Japan) connected to the soundcard. The loudspeakers’ frequency response and output level was measured at the moth’s position using a calibrated measuring microphone (40BF, with pre-amplifier 26AC and Power Module 12AA, GRAS Sound & Vibration, Holte, Denmark). Two infrared video cameras (Flea 3, FLIR Integrated Imaging Solutions, Richmond, Canada, with HF12.5HA-1B lenses, Fujinon, Tokyo, Japan) at 30 cm distance below and in front of the moth recorded its behaviour, illuminated by four infrared lights (850 nm, Mini IR Illuminator TV6700, ABUS Security-Center, Affing, Germany) installed in the corners around the frontal camera. Stimulus presentation and data acquisition of flight recorder and cameras was controlled with custom MATLAB code (The Mathworks Inc., Natick, Massachusetts, USA).

**Figure 1:**
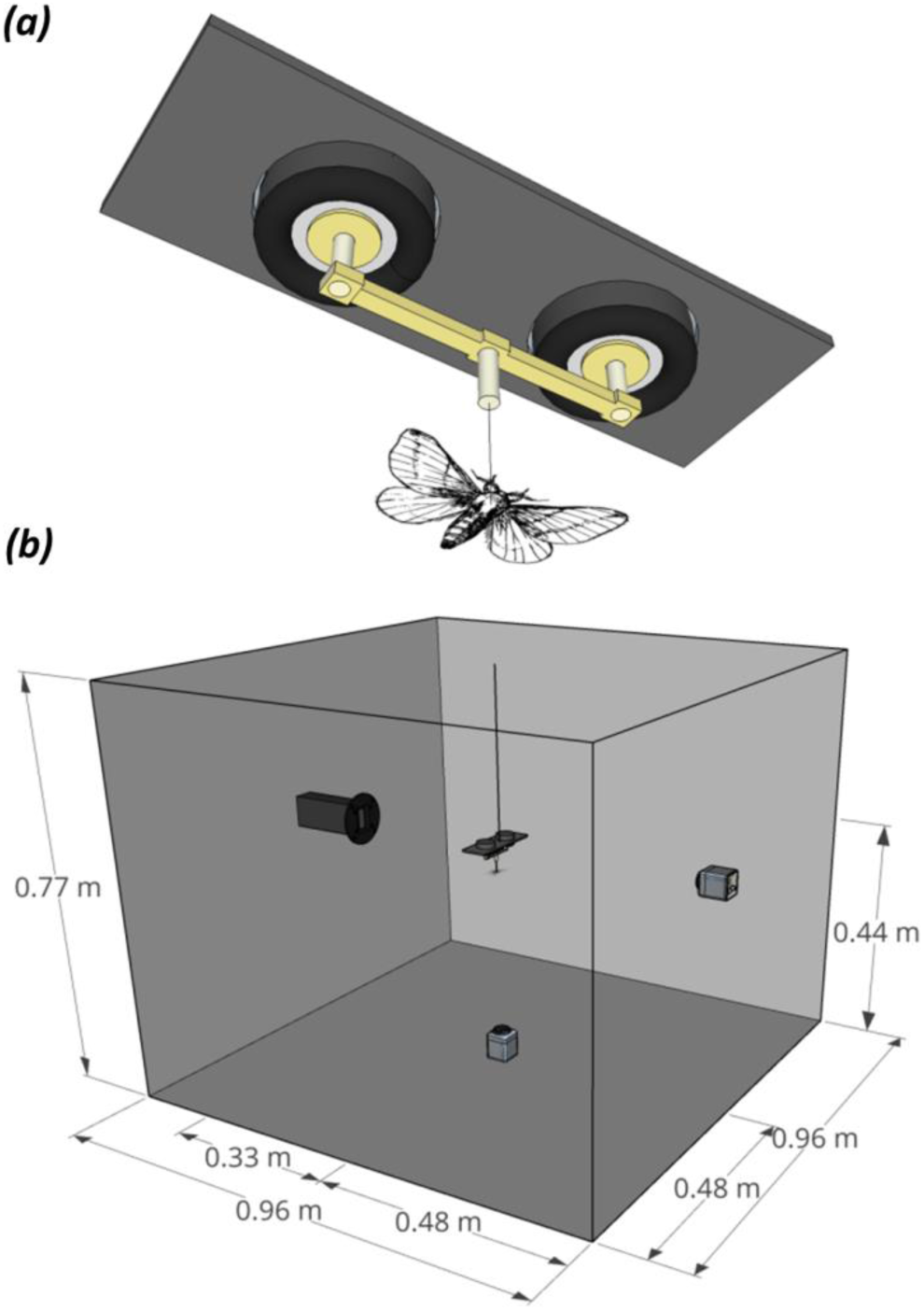
flight recorder system for recording flight strength. **(*a*)** Flight recorder consisting of two speakers connected by a wooden connector, which held the moth via a needle glued to the moth’s thorax. **(*b*)** The flight recorder was mounted in the centre of an anechoic chamber, which also hold an ultrasonic speaker (left) to present stimuli and two IR-cameras (right and bottom). Moths were orientated to face away from the speaker.

### Stimulus

Last ditch flight behaviour in many moths is argued to be elicited by activation of the auditory receptor neuron A2 (Gordon & Ter Hofstede, 2018; Madsen & Miller, 1987; Kenneth D. Roeder, 1974). A2 sensitivity depends on frequency, having highest sensitivity between 15 to 60 kHz (Gordon & Ter Hofstede, 2018; Madsen & Miller, 1987; Surlykke, 2003; Ter Hofstede, Goerlitz, Ratcliffe, Holderied, & Surlykke, 2013; Waters & Jones, 1996; Zha, Chen, & Lei, 2009). We therefore designed a stimulus to elicit last ditch flight behaviour and mimicking an attacking bat (Schnitzler & Kalko, 2001; Schnitzler, Moss, & Denzinger, 2003; Skiba, 2014), consisting of 120 pure tones at 35 kHz, each having 4 ms duration plus 0.5 ms raised-cosine-ramps and 25 ms pulse interval (PI), resulting in a total length of 3 s. Sound pressure level at the moth was 80 dB SPL RMS re. 20 μPa, i.e., about 8 to 17 dB above A2 threshold depending on moth species (Gordon & Ter Hofstede, 2018; Surlykke, 2003; Ter Hofstede et al., 2013; Zha et al., 2009). We recorded moth flight strength from 2 sec before until 1 sec after stimulus presentation, resulting in a 6 s-long recording in total. Stimuli were presented to non-flying and flying moths.

### Moth species

We conducted the experiment at the Siemers Bat Research Station in Tabachka, N-Bulgaria, between 17 June and 22 August 2017. We caught moths with two light traps (Sylvania, blacklight, F15W/350BL-T8, USA) between dusk and midnight on a plateau with wild meadow and some scattered bushes. Since we caught all moths of eight different species at this one site, all species use a similar habitat and activity period. All of those moth species thus experience a similar predation threat by bats, and each predatory bat hunting in this habitat will encounter all of those moth species. Caught moths were individually kept in Falcon tubes until being tested on the same night. Species were identified using Steiner et al. 2014 (Steiner, Ratzel, Top-Jensen, & Fibiger, 2014). We selected moth species based on availability and on the absence of any other known anti-predator strategy in that species, such as chemical defence or jamming. We tested 172 individuals of eight different eared species and two families: *Amphipyra pyramidae, Helicoverpa armigera, Heliothis adaucta, Noctua comes, Noctua fimbriata, Noctua janthe* and *Xestia c-nigrum*, which all belong to the family of Noctuidae, and *Deilephia porcellus* belonging to the family of Sphingidae. To attach the insect-pin to a moth, we placed individual moths on a piece of foam, held it in place with a soft, coarse-meshed plastic grid, removed the scales on the thorax gently with a scalpel, and glued the flat head of an insect needle to the thorax using cyanoacrylate glue. As soon as the glue set, the needle was inserted into the flight recorder with the moth facing away from the loudspeaker.

### Surface area measurement

We measured the surface area of 130 moths, mainly (72.7 %) overlapping with tested individuals (see electronic supplementary material, Table S1). All individuals were deep frozen for at least 24 hours and then fixed on a sheet of squared paper with completely spread wings to ensure maximum surface area. Photos were taken from fixed distance and surface area was measured using Image J (National Institute of Health, Bethesda, USA). We converted photos of moths into 8bit-black-and-white-images, manually adjusted their intensity threshold to detect moth area. We used the automatic outline detection to detect the moth’s outline and calculated the area within the outline. Measured surface area of individuals covered a range from 343 to 1008 mm^2^ (see electronic supplementary material, Fig. S1). We assigned the species’ mean value to individuals without size measurements for subsequent statistical analyses. This does not affect the potential correlation between size and flight strength, but makes its detection more difficult due to reduced variation in size, making our analysis more conservative.

### Analysis of flight strength

The flight movement of the tethered moths moved the two loudspeaker membranes and thus generated voltage fluctuations, which we recorded from 2 sec before until 1 sec after presentation of the 3-sec-long stimulus. For further analysis, we only analysed the time period around stimulus onset, from 1 s before to 1 s after stimulus onset. We calculated the root-mean-square (RMS) of the recorded voltage per 100 ms bins, resulting in 20 measurements per speaker over the analysed duration of 2 s. We express the measurements in dB FS RMS, i.e., as negative values on a logarithmic scale relative to the highest recordable voltage of the flight recorder (= 0 dB FS; FS: full scale), and as the mean of both loudspeakers. We will refer to this value as flight strength, since low to high values correspond to non-flying moths to different degrees of flight activity in flying moths. Since we recorded nine and three individuals two and three times, respectively, we used for each individual only the first recording for analysis.

We performed a Principal Component Analysis (PCA) to identify different types of reaction in response to simulated bat calls, such as onset/cessation of flight or increase/decrease in flight strength with stimulus onset. Classically, PCA is used to reduce the number of correlated explanatory variables to fewer uncorrelated variables, the so-called Principle Components (PCs) that have the most explanatory power. In our case, we used the PCA to reduce the number of 20 correlated flight strength measurements collected over 20 time bins to a smaller number of PCs that reflect the most common types of reaction. For each PC, we obtained one loading per time bin (i.e., 20 loadings in total). Combining loadings with PC-scores reconstructs the original flight strength data; comparing loadings over time bins therefore revealed patterns of changing flight strength over time.

Some moths flew irregularly or stopped flying after some time. We thus split our dataset into moths that were flying before stimulus onset (“active moths”) and those that were not flying (“inactive moths”). Initial analysis showed that flight strengths in non-flying moths was −83 dB FS RMS, while flying moths had higher values. Hence, we set the threshold between active and non-active conservatively to −80 dB FS RMS and analysed flight strength in the bins at 1.0 – 0.9 s and 0.1 – 0 s before stimuli onset. If flight strength was above threshold in both bins, an individual was defined as “active” (N=92; Table S1), if flight strength was below threshold in both bins, it was defined as “inactive” (N=74; Table S1). Six individuals had flight strengths once above and once below threshold in those bins and were excluded (Table S1). We used linear models to fit PCs as a function of the fixed effects species (categorical) and surface area (continuous; R version 3.3.2, R Foundation for Statistical Computing, Vienna, Austria; RStudio, version 1.1.463, RStudio, Bosten, USA). We tested for a significant effect of factors on PC-values using likelihood ratio tests to compare the full model to the model excluding a factor.

## RESULTS

In total, we tested the flight behaviour of 172 individuals of eight eared moth species. 92 individuals were flying before stimulus onset (“active moths”), while 74 were non-flying before stimulus onset (“non-active moths”; Table S1). Since we were interested to study evasive flight in flying moths, we focus in the following on the active moths. Sample size varied between species as this depended on moth availability, ranging from 33 (*H*. *armigera*) to two individuals (*H. adaucta, D. porcellus*).

Reactions to the acoustic stimulus were mostly consistent within a species, but differed between species (Fig. 2). We found three main reaction types in active moths, either constant flight strength over time, or increasing or decreasing flight strength after stimulus onset. In addition, both the median flight strength and its inter-individual variation differed between moth species. For example, *H*. *armigera* and *N. fimbriata* both showed fairly constant flight strength over time, but differed in their median and inter-individual variation. Flight strength of the 33 individuals of *H*. *armigera* ranged from −80 to −50 dB FS, with a median around −63 dB FS, while all 12 individuals of *N. fimbriata* had a constant and high flight strength around −50 dB FS. *X. c-nigrum* and *N. janthe* increased their flight strength after stimulus onset, with *N. janthe* (N=32) also showing a clear reduction in inter-individual variation after stimulus onset. *H. adaucta, D. porcellus* and *N. comes* all decreased their flight strength, with additional variation in overall flight strength, timing and exact temporal pattern of the change. The three individuals of *A. pyramidae* showed a mix constant, increasing and decreasing flight strength.

**Figure 2:**
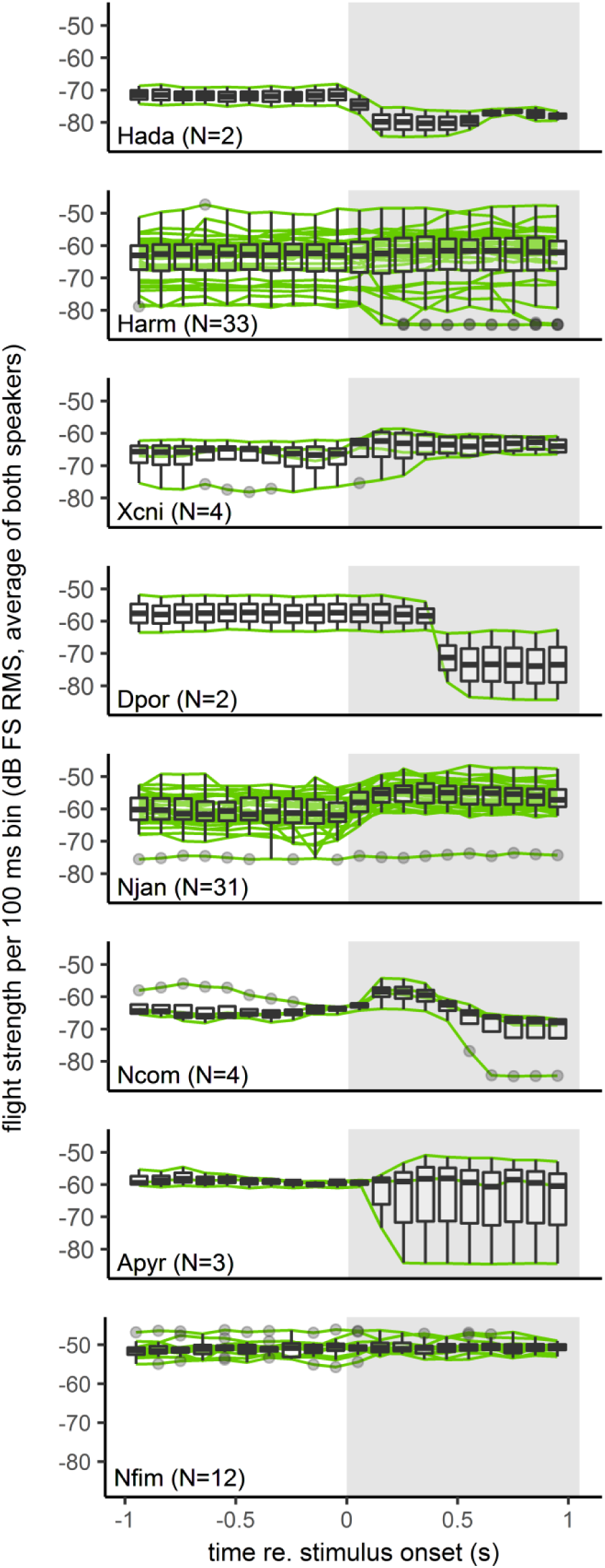
Flight strength per 100 ms bin for active moths from 1 s before to 1s after the stimulus onset for 8 moth species. Species are ordered with decreasing surface area from top to bottom. Green lines are individual data; boxplots show median, quartiles, whiskers (1.5 × inter-quartile range) and outliers beyond the interquartile range. Stimulus presentation is indicated by the grey shading. Species abbreviations: Hada = *Heliothis adaucta*, Harm = *Helicoverpa armigera*, Xcni= *Xestia c-nigrum*, Dpor = *Deilephia porcellus*, Njan = *Noctua janthe*, Ncom= *Noctua comes*, Apyr= *Amphipyra pyramidae*, Nfim = *Noctua fimbriata*.

We used a PCA to reduce the temporal correlation in flight strength measurements and to obtain behavioural categories for testing the above observations. The first two components of the PCA of active moths explain 95.7% of the overall variation in flight strength (Fig. 3a, PC1: 82.6%, PC2: 13.1%), with overlapping clusters in PC1 and PC2 scores between species (Fig. 3b). Loadings of these two components (electronic supplementary material, Table S2) match the observed reaction types towards the stimulus. Loadings for PC1 are almost constant over time. Hence, PC1 scores describe the general flight strength of an individual (Fig. 3c). Loadings for PC2 invert their sign over time (from negative before stimulus onset to positive after stimulus onset) and therefore describe the temporal pattern of flight strength, which can be either increasing (for positive individual PC2 scores), constant (for PC2 scores close to Zero) or decreasing (for negative individual PC2 scores; Fig. 3d). Hence, PC2 scores describe the reaction type of an individual.

**Figure 3:**
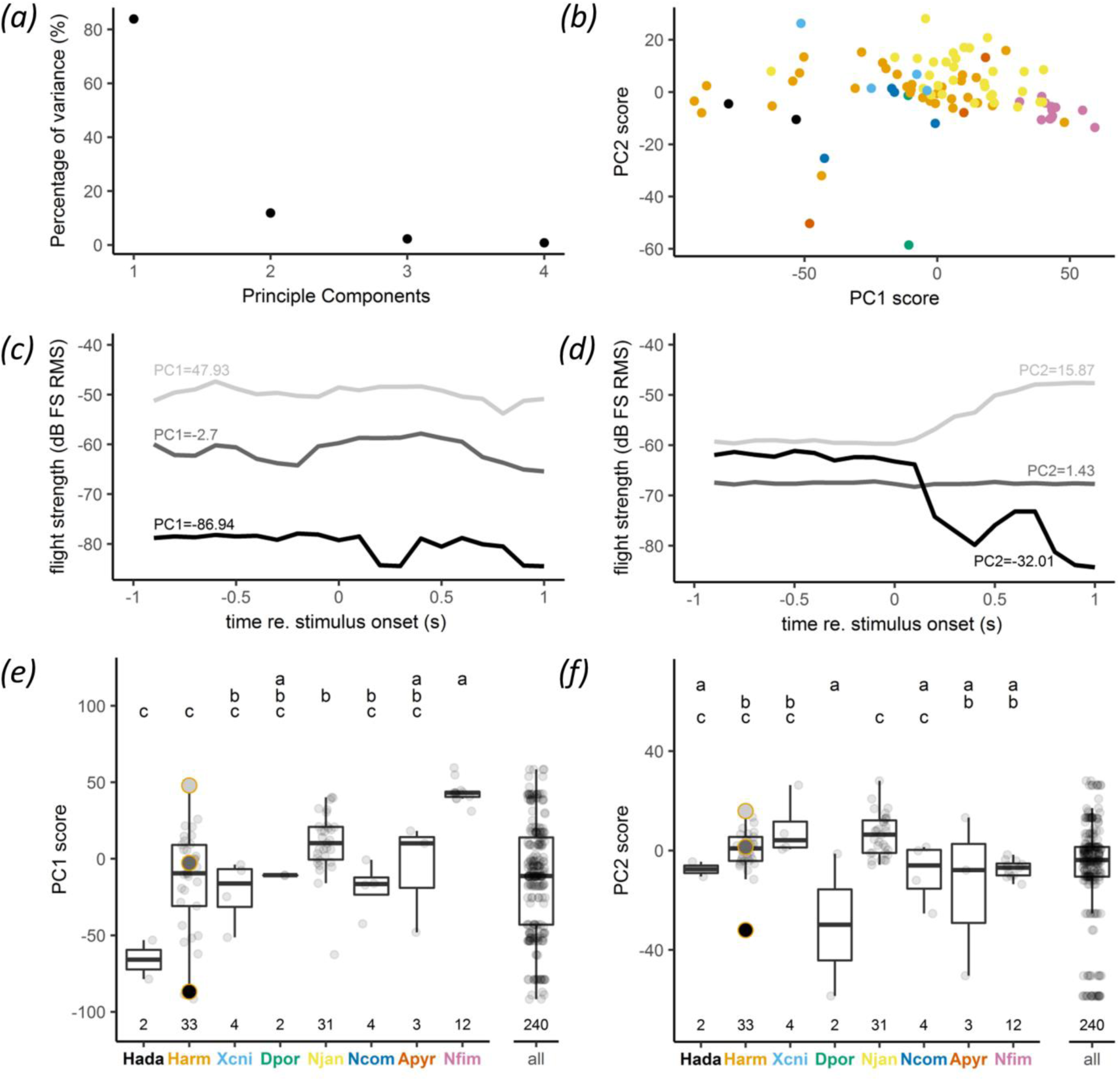
Principle components of moths flying before stimulus onset (“active” moths). (*a*) Percentage of variance explained by the first four principle components. (*b*) PC1 scores as a function of PC2 scores; for species colour code see panels e) and g). (*c, d*) Example flight behaviours for high (light grey), medium (dark grey) and low (black) values for PC1 and PC2. (*e, f*) PC1 and PC2 scores for each species and for the whole population (“all”), which was created by randomly sampling 30 values from each species. Species are ordered with increasing surface area from left to right. Letters above boxes indicate significant differences. Circled data points refer to examples shown above. Boxplots show median, quartiles, whiskers (1.5 × inter-quartile range) and outliers beyond the whiskers.

We found significant species-specific differences in the PC1 and PC2 scores, confirming species-specific flight strength and reaction types in response to acoustic stimuli (Figs 3e-f). While the whole population of all tested moth species shows a large variation in PC1 and PC2 scores, each species only clusters within a smaller range of PC1 and PC2 scores, respectively (Figs 3e-f, compare ‘all’ versus each species). PC1 scores were significantly affected by species (p< 0.001), but not by surface area (p= 0.64). The PC1 score captures the species-specific median and variation in flight strength observed before. For example, *N. fimbriata* having high and constant flight strength also has high PC1 scores with little variation. *H*. *armigera*, having intermediate flight strength with high variation, also has intermediate PC1 scores with high variation; and *H. adaucta* shows both low flight strength and low PC1 scores. PC2 scores were significantly affected by species (p<0.001), but not by surface area (p= 0.34). The PC2 score captures the species-specific reaction type, i.e., the change in flight strength over time as observed before. For example, *H*. *armigera* and *N. fimbriata* have fairly constant flight strength over time, and correspondingly have PC2 scores close to Zero. Correspondingly, positive (*X. c-nigrum, N. janthe*) and negative (*H. adaucta, D. porcellus, N. comes*) PC2 scores capture increasing and decreasing flight strength over time, respectively.

In addition to the 92 active moths that were flying at stimulus onset, we also tested 74 non-active moths that were motionless at stimulus onset (Fig. S2, S3). Despite the very different initial state of the moths (flying vs. non-flying), we can see similar behavioural changes. Particularly for *N. janthe*, most of the 57 non-active individuals started to fly after stimulus onset, matching the increased flight strength observed in active moths.

## DISCUSSION

Using standardized measures in a novel flight recorder, we show for the first time species-specific reactions in eared moths in response to the same bat-like sounds, and thus inter-species differences in evasive flight strategies within a sympatric moth community. These clear species-specific reactions are particularly remarkable given our rather simple measured variable and the artificial tethered flight which limits moths in their flight behaviour and probably affects their motor-sensory feedback loop (Taylor, 2001). Although free-flight experiments will be required to link vertical flight strength in tethered flight to actual three-dimensional flight trajectories, free-flight studies showed a clear change of kinematic parameters during last-ditch evasive flight (Corcoran & Conner, 2016, 2017). Our observations are further supported because the reaction of a given species was independent of whether individuals were flying or not at stimulus onset. This suggests that last-ditch evasive flight is to some extent hardwired and can be elicited by the appropriate acoustic input. In addition, however, a substantial amount of variation in flight strength also existed between individuals in some species. Whether some moths furthermore show variation within individuals between subsequent executions of evasive flight is yet unknown. In summary, our data supports the hypothesis of escape-tactic diversity in moths. An echolocating bat preying on a multi-species prey community with prey-species-specific differences in evasive flight faces larger variation and unpredictability than would be generated by any single species. Species-specific differences in evasive flight within prey communities thus likely provide increased protection against predators for each individual in the community, particularly if the predator does not discriminate between prey species. In this case, it is not even necessary that prey individuals form actual groups of multiple individuals that the predator encounters simultaneously. It is sufficient that different species, which are attacked but not discriminated by the same predator, react in different ways to prevent the predator from predicting the type of evasive action for each independent attack. As we caught all moths in the same field site, a predatory bat has a high likelihood of encountering different species while hunting in this habitat.

Our data of eight moth species suggest two key components of last-ditch evasive flight: overall flight strength and temporal reaction type, with each species showing its unique combination and therefore filling its own ecological niche. We could only test a large number of individuals in three species (*Noctua fimbriata, Helicoverpa armigera, Noctua janthe*). Even though all three belong to the family Noctuidae, they exhibited different strategies for evasive flight. *N. fimbriata* flew constantly strong without intraspecific variation and no temporal change in response to acoustic stimulation. Similarly, *H. armigera* flew constantly with intermediate flight strength and no temporal change in response to acoustic stimulation, yet showed strong intra-specific variation. *N. janthe* also flew with intermediate flight strength with intra-specific variation, yet all individuals increased their flight strength after acoustic stimulation. What might our observations in the flight recorder mean under real world free-flight conditions? Of the three species with high sample size, *N. janthe* was the only one with a clear change in flight strength with stimulus onset. It increased its flight strengths after stimulus onset and reached maximum flight strength within 200-300 ms after stimulus onset, corresponding to 8-12 pulses of our acoustic pulse train. Arguably, this increase in flight strength corresponds to a certain kind of last ditch flight behaviour under natural free-flight conditions. *H. armigera* did not change its flight strengths, yet showed a high variability in flight strength between individuals. This inter-individual variability might relate to a generally variable flight behaviour between individuals that could already function as general anti-predator strategy. *N. fimbriata* showed the strongest and the most uniform flight strength of all tested species, without a reaction to the acoustic stimulus. The high flight strength of this species might indicate that it is a fast flyer and thus difficult to catch. This is supported by the large size of *N. fimbriata* (932.6 ± 54.8 mm^2^, mean±std), which is positively correlated with acceleration in Lepidoptera (Berwaerts et al., 2002). The remaining moth species with lower sample size showed further variation in general flight strength and its temporal pattern, suggesting that they exhibit different flight trajectories in response to acoustic stimulation. For example, *Noctua comes* showed increasing and decreasing flight strength, which might correlate with acceleration and subsequence deceleration. *Heliothis adaucta* and *Deilephia porcellus* both reduced flight strength, however with different temporal patterns: *H. adaucta* decreased flight strength already within the first 100 ms and reached its minimum within 200 ms, while *D. porcellus* decreased flight strength only after 400 ms. Both patterns might represent different types of (power) dives in response to an attacking bat. Interestingly, *D. porcellus* flew with higher flight strength before stimulus onset than *H. adaucta*, which might represent a higher flight speed, allowing it to react later to an attacking bat.

As *N. fimbriata* and *H. armigera* did not change their flight strength in response to our stimulus, consisting of 35 kHz pure tones at 80 dB SPL RMS, it is possible that our stimulus was inaudible for those species, or audible yet too faint to trigger evasive flight or to be perceived as sufficiently high predation risk. Although neuronal audiograms of multiple species suggest that our stimulus is above the threshold of the A2-cell of moths (Gordon & Ter Hofstede, 2018; Surlykke, 2003; Ter Hofstede et al., 2013; Zha et al., 2009), little is known about how neuronal activity translates into evasive flight. Behavioural thresholds are generally higher than neuronal thresholds, although the exact differences and potential variation between species are mostly unknown (for discussion, see (Lewanzik & Goerlitz, 2017)). Variation in the translation from neuronal activity to evasive flight might even add additional unpredictability to the evasive flight of moths. Lastly, additional anti-predator strategies could reduce the need for evasive flight. Although we caught all moths in the same open-field habitat, moths might still possess species-specific differences in flight behaviour. For example, flying closer to the ground or vegetation could be a potential anti-predator strategy, as close-by background structures impairs bats’ capture success due to sensory and motor constraints (Siemers & Schnitzler, 2004).

Although size affects flight capabilities, we did not detect an effect of size on flight strength (PC1) or main temporal reaction type (PC2). While a direct influence of size on temporal reaction type is not obvious, we would have expected to find a positive correlation between moth size and flight strength. The lack of this correlation might be due to the small number of individuals for some species, or too few species tested altogether; or it might be a true effect. The lack of this correlation might have been driven by the benefits of increased unpredictability, reducing size-dependent constraints on flight strength.

## CONCLUSION

Our data provide novel insights into the function and evolution of defensive strategies in mixed-species prey communities. We show that a basic measure, such as vertical flight strength, can reveal both stereotypy and variability in escape strategies within and between species. We show that evasive flight in moths is more variable on the community level than within any single species, confirming the escape-tactic diversity hypothesis for eared moths. This inter-specific variability adds to the total unpredictability of evasive flight that a predator experiences, and suggests an emergent benefit of overlapping in time and space for prey animals.

## DATA AVAILABILITY

Data and code will be made available at DRYAD

## AUTHORS’ CONTRIBUTIONS

- HRG conceived, designed and supervised the study
- TH contributed to study design and collected the data
- TH analysed the data with input of HRG
- TH and HRG wrote the manuscript

## ACKNOWLEDGEMENTS

We thank Erich Koch for help with setup development and Felix Hartl for help with the light traps. We are grateful to Boyan Zlatkov for the assistance with moth identification. We thank the Max Planck Institute for Ornithology Seewiesen for providing excellent infrastructure and support, and the IMPRS for Organismal Biology for support and funding. Further, we thank Martin Yordanov Georgiev for help with data collection, Fränzi Korner-Nievergelt and Tobias Roth for statistical advice, and Sue Anne Zollinger for comments on a previous version this manuscript.

## References

Agee, H. R. (1969). Response of flying bollworm moths and other tympanate moths to pulsed ultrasound. Ann Entomol Soc Am, 62(4), 801–807. https://doi.org/10.1093/aesa/62.4.801

Anthony, E. L. P., & Kunz, T. H. (1977). Feeding strategies of the little brown bat, *Myotis Lucifugus*, in southern New Hampshire. Ecology, 58(4), 775–786. https://doi.org/10.2307/1936213

Berwaerts, K., Van Dyck, H., & Aerts, P. (2002). Does flight morphology relate to flight performance? An experimental test with the butterfly *Pararge aegeria*. Funct Ecol, 16(4), 484–491. https://doi.org/10.1046/j.1365-2435.2002.00650.x

Bogdanowicz, W., Fenton, M. B., & Daleszczyk, K. (1999). The relationships between echolocation calls, morphology and diet in insectivorous bats. J Zool, 247(3), 381–393. https://doi.org/10.1111/j.1469-7998.1999.tb01001.x

Conrad, K. F., Woiwod, I. P., Parsons, M., Fox, R., & Warren, M. S. (2004). Long-term population trends in widespread British moths. J Insect Conserv, 8(2), 119–136. https://doi.org/10.1023/B:JICO.0000045810.36433.c6

Corcoran, A. J., & Conner, W. E. (2012). Sonar jamming in the field: effectiveness and behavior of unique prey defense. J Exp Biol, 215(Pt 24), 4278–4287. https://doi.org/10.1242/jeb.076943

Corcoran, A. J., & Conner, W. E. (2016). How moths escape bats: predicting outcomes of predator-prey interactions. J Exp Biol, jeb.137638. https://doi.org/10.1242/jeb.137638

Corcoran, A. J., & Conner, W. E. (2017). Predator counteradaptations: stealth echolocation overcomes insect sonar-jamming and evasive-manoeuvring defences. Animal Behav, 132, 291–301. https://doi.org/10.1016/J.ANBEHAV.2017.08.018

Denzinger, A., & Schnitzler, H.-U. (2013). Bat guilds, a concept to classify the highly diverse foraging and echolocation behaviors of microchiropteran bats. Front Psychol, 4, 1–15.

Dudley, R. (2002). Mechanisms and implications of animal flight maneuverability. Integr Comp Biol, 42(1), 135–140. https://doi.org/10.1093/icb/42.1.135

Fenton, M. B., Portfors, C. V., Rautenbach, I. L., & Waterman, J. M. (1998). Compromises: sound frequencies used in echolocation by aerial-feeding bats. Can J Zool, 76(6), 1174–1182. https://doi.org/10.1139/z98-043

Findley, J. S., & Black, H. (1983). Morphological and dietary structuring of a Zambian insectivorous bat community. Ecology, 64(4), 625–630. https://doi.org/10.2307/1937180

Fullard, J. H., Muma, K. E., & Dawson, J. W. (2003). Quantifying an anti-bat flight response by eared moths. Can J Zool, 81(3), 395–399. https://doi.org/10.1139/z03-019

Göpfert, M. C., & Wasserthal, L. T. (1999). Auditory sensory cells in hawkmoths: identification, physiology and structure. J Exp Biol, 202, 1579–1587.

Gordon, S. D., & Ter Hofstede, H. M. (2018). The influence of bat echolocation call duration and timing on auditory encoding of predator distance in noctuoid moths. J Exp Biol, 221(6), jeb171561. https://doi.org/10.1242/jeb.171561

Hölker, F., Dörner, H., Schulze, T., Haertel-Borer, S. S., Peacor, S. D., & Mehner, T. (2007). Species-specific responses of planktivorous fish to the introduction of a new piscivore: implications for prey fitness. Freshw Biol, 52(9), 1793–1806. https://doi.org/10.1111/j.1365-2427.2007.01810.x

Howland, H. C. (1974). Optimal strategies for predator avoidance: The relative importance of speed and manoeuvrability. J Theor Biol, 47(2), 333–350. https://doi.org/10.1016/0022-5193(74)90202-1

Humphries, D. A., & Driver, P. M. (1967). Erratic display as a device against predators. Science, 156(3783), 1767–1768. https://doi.org/10.1126/SCIENCE.156.3783.1767

Humphries, D. A., & Driver, P. M. (1970). Protean defense by prey animals. Oecologia, 5, 285–302. https://doi.org/10.1007/BF00815496

Lewanzik, D., & Goerlitz, H. R. (2017). Continued source level reduction during attack in the low- amplitude bat *Barbastella barbastellus* prevents moth evasive flight. Funct Ecol, 32(5), 1251–1261. https://doi.org/10.1111/1365-2435.13073

Madsen, B. M., & Miller, L. A. (1987). Auditory input to motor neurons of the dorsal longitudinal flight muscles in a noctuid moth (*Barathra brassicae L.*). J Comp Physiol A, 160(1), 23–31. https://doi.org/10.1007/BF00613438

Nakano, R., Ihara, F., Mishiro, K., Toyama, M., & Toda, S. (2015). High duty cycle pulses suppress orientation flights of crambid moths. J Insect Physiol, 83, 15–21. https://doi.org/10.1016/j.jinsphys.2015.11.004

Randall, J. A., Hatch, S. M., & Hekkala, E. R. (1995). Inter-specific variation in anti-predator behavior in sympatric species of kangaroo rat. Behav Ecol Sociobiol, 36(4), 243–250. https://doi.org/10.1007/BF00165833

Roeder, K D. (1966). A differential anemometer for measuring the turning tendency of insects in stationary flight. Science, 153(3744), 1634–1636.

Roeder, Kenneth D. (1962). The behaviour of free flying moths in the presence of artificial ultrasonic pulses. Anim Behav, 10(3–4), 300–304. https://doi.org/10.1016/0003-3472(62)90053-2

Roeder, Kenneth D. (1974). Responses of the less sensitive acoustic sense cells in the tympanic organs of some noctuid and geometrid moths. J Insect Physiol, 20(1), 55–66. https://doi.org/10.1016/0022-1910(74)90123-1

Rydell, J., Roininen, H., & Philip, K. W. (2000). Persistence of bat defence reactions in high arctic moths (Lepidoptera). Proc Royal Soc B, 267(1443), 553–557. https://doi.org/10.1098/rspb.2000.1036

Rydell, J., Skals, N., Surlykke, A., & Svensson, M. (1997). Hearing and bat defence in geometrid winter moths. Proc Biol Sci, 264(1378), 83–88. https://doi.org/10.1098/rspb.1997.0012

Rydell, Jens, Jones, G., & Waters, D. A. (1995). Echolocating bats and hearing moths: Who are the winners? Oikos, 73, 419–424. https://doi.org/Doi10.2307/3545970

Scalercio, S., Infusino, M., & Woiwod, I. P. (2008). Optimising the sampling window for moth indicator communities. J Insect Conserv, 13(6), 583. https://doi.org/10.1007/s10841-008-9206-x

Schall, J. J., & Pianka, E. R. (1985). Evolution of escape behavior diversity. Am Nat, 115(4), 551–566. http://dx.doi.org/10.2307/2460484

Schnitzler, H.-U., & Kalko, E. K. V. (2001). Echolocation by insect-eating bats. BioScience, 51(7), 557–569. https://doi.org/10.1641/0006-3568(2001)051[0557:EBIEB]2.0.CO;2

Schnitzler, H.-U., Moss, C. F., & Denzinger, A. (2003). From spatial orientation to food acquisition in echolocating bats. Trends Ecol Evol, 18, 386–394. https://doi.org/10.1016/S0169-5347(03)00185-X

Siemers, B. M., & Schnitzler, H.-U. (2004). Echolocation signals reflect niche differentiation in five sympatric congeneric bat species. Nature, 429, 657–661. https://doi.org/10.1038/nature02547

Skiba, R. (2014). Europäische Fledermäuse (2. Edition). Magdeburg: VerlagsKG Wolf.

Steiner, A., Ratzel, U., Top-Jensen, M., & Fibiger, M. (2014). Die Nachtfalter Deutschlands. Ein Feldführer. Oestermarie, Denmark: Bugbook Publishing.

Summerville, K. S., & Crist, T. O. (2004). Contrasting effects of habitat quantity and quality on moth communities in fragmented landscapes. Ecography, 27(1), 3–12. https://doi.org/10.1111/j.0906-7590.2004.03664.x

Surlykke, A. (2003). Hearing in hooktip moths (Drepanidae: Lepidoptera). J Exp Biol, 206(15), 2653–2663. https://doi.org/10.1242/jeb.00469

Svensson, M., Rydell, J., & Brown, R. (1999). Bat predation and flight timing of winter moths, *Epirrita* and *Operophtera* species (Lepidoptera, Geomentridae). Oikos, 84(2), 193–198. https://doi.org/10.2307/3546713

Taylor, G. K. (2001). Mechanics and aerodynamics of insect flight control. Biol Rev, 76, 449–471. https://doi.org/10.1017\S1464793101005759

Ter Hofstede, H. M., Goerlitz, H. R., Ratcliffe, J. M., Holderied, M. W., & Surlykke, A. (2013). The simple ears of noctuoid moths are tuned to the calls of their sympatric bat community. J Exp Biol, 216(21), 3954–3962. https://doi.org/10.1242/jeb.093294

Ter Hofstede, H. M., & Ratcliffe, J. M. (2016). Evolutionary escalation: the bat-moth arms race. J Exp Biol, 219(Pt 11), 1589–1602. https://doi.org/10.1242/jeb.086686

Treat, A. E. (1955). The response to sound in certain Lepidoptera. Ann Entomol Soc Am, 48(4), 272–284. https://doi.org/10.1093/aesa/48.4.272

Truxa, C., & Fiedler, K. (2012). Down in the flood? How moth communities are shaped in temperate floodplain forests. Insect Conserv Divers, 5(5), 389–397. https://doi.org/10.1111/j.1752-4598.2011.00177.x

Waters, D. A. (2003). Bats and moths: What is there left to learn? Physiol Entomol, 28(4), 237–250. https://doi.org/10.1111/j.1365-3032.2003.00355.x

Waters, D. A., & Jones, G. (1996). The peripheral auditory characteristics of noctuid moths: Responses to the search-phase echolocation calls of bats. J Exp Biol, 199(4), 847–856.

Wilson, A. M., Hubel, T. Y., Wilshin, S. D., Lowe, J. C., Lorenc, M., Dewhirst, O. P., … West, T. G. (2018). Biomechanics of predator–prey arms race in lion, zebra, cheetah and impala. Nature, 554, 183–188. https://doi.org/10.1038/nature25479

Wohlfahrt, B., Mikolajewski, D. J., Joop, G., & Suhling, F. (2006). Are behavioural traits in prey sensitive to the risk imposed by predatory fish? Freshw Biol, 51(1), 76–84. https://doi.org/10.1111/j.1365-2427.2005.01475.x

Wolf, N. G. (1985). Odd fish abandon mixed-species groups when threatened. Behav Ecol Sociobiol, 17(1), 47–52. https://doi.org/10.1007/BF00299428

Zha, Y., Chen, Q., & Lei, C. (2009). Ultrasonic hearing in moths. Ann Soc Entomol Fr, 45(2), 145–156. https://doi.org/10.1080/00379271.2009.10697598

