## Supplementary material for "Species-specific strategies increase unpredictability of escape flight in eared moths" (from another version of this preprint: version 1, 2018-12-03)

### Supplementary information

| | behavioural test | | | | size measurements | | | |
|---|---|---|---|---|---|---|---|---|
| | active | non-active | excluded | total tested | active | non-active | measured and tested | total measured |
| **Hada** | 2 | 0 | 1 | 3 | 2 | 0 | 3 | 3 |
| **Harm** | 33 | 2 | 1 | 36 | 20 | 1 | 21 | 21 |
| **Xcni** | 4 | 6 | 0 | 10 | 2 | 2 | 4 | 4 |
| **Dpor** | 2 | 4 | 0 | 6 | 2 | 1 | 3 | 3 |
| **Njan** | 32 | 57 | 2 | 91 | 26 | 45 | 73 | 76 |
| **Ncom** | 4 | 2 | 1 | 7 | 3 | 1 | 4 | 5 |
| **Apyr** | 3 | 1 | 0 | 4 | 3 | 1 | 4 | 4 |
| **Nfim** | 12 | 2 | 1 | 15 | 11 | 1 | 13 | 14 |
| | 92 | 74 | 6 | 172 | 69 | 52 | 125 | 130 |

**Table S1. Sample sizes of moths used for behavioural tests and size measurements.** For behavioural test data splits into individuals that were active or not active before the stimulus onset. Single individuals that did not fulfil criteria for active or non-active were excluded. Size measurements were taken from individuals that were used in behavioural tests, but also from additional individuals. Numbers for active and non-active indicate number of individuals with individual size measurements, for the active and non-active dataset respectively. All other individuals were assigned with the species-specific mean of size measurement.

| ACTIVE | | | | | | | | | | | | | | | | | | | | |
|---|---|---|---|---|---|---|---|---|---|---|---|---|---|---|---|---|---|---|---|---|
| **bins** | **PC1** | **PC2** | **PC3** | **PC4** | **PC5** | **PC6** | **PC7** | **PC8** | **PC9** | **PC10** | **PC11** | **PC12** | **PC13** | **PC14** | **PC15** | **PC16** | **PC17** | **PC18** | **PC19** | **PC20** |
| 1 | 0.19 | -0.20 | 0.00 | -0.27 | 0.00 | 0.00 | 0.00 | -0.23 | -0.31 | 0.30 | 0.38 | -0.14 | -0.12 | 0.11 | -0.36 | 0.23 | -0.33 | 0.34 | 0.00 | 0.00 |
| 2 | 0.20 | -0.21 | 0.00 | -0.31 | 0.00 | 0.00 | 0.29 | -0.17 | 0.00 | 0.26 | -0.12 | 0.21 | 0.16 | 0.22 | 0.24 | 0.26 | 0.47 | -0.18 | 0.29 | 0.17 |
| 3 | 0.20 | -0.23 | -0.10 | -0.33 | 0.00 | -0.10 | 0.17 | -0.11 | 0.00 | 0.17 | -0.27 | 0.24 | 0.00 | -0.32 | 0.20 | -0.47 | -0.29 | 0.00 | -0.35 | 0.00 |
| 4 | 0.20 | -0.22 | -0.13 | -0.30 | 0.00 | -0.21 | 0.00 | 0.23 | 0.22 | -0.54 | -0.19 | 0.00 | -0.30 | -0.15 | -0.39 | 0.24 | 0.00 | 0.00 | 0.00 | 0.00 |
| 5 | 0.20 | -0.23 | 0.00 | -0.28 | 0.00 | 0.00 | -0.25 | 0.29 | 0.23 | -0.15 | 0.27 | -0.35 | 0.51 | 0.28 | 0.13 | -0.20 | 0.00 | 0.00 | 0.00 | 0.00 |
| 6 | 0.20 | -0.21 | 0.00 | 0.00 | 0.00 | 0.00 | -0.42 | 0.00 | -0.17 | 0.00 | 0.00 | -0.29 | -0.46 | -0.14 | 0.25 | 0.00 | 0.21 | -0.18 | 0.18 | -0.45 |
| 7 | 0.20 | -0.22 | 0.00 | 0.20 | 0.00 | 0.00 | -0.47 | 0.00 | -0.13 | 0.00 | -0.17 | 0.00 | 0.00 | -0.16 | 0.00 | 0.22 | 0.00 | 0.00 | -0.20 | 0.67 |
| 8 | 0.20 | -0.23 | 0.00 | 0.37 | 0.00 | 0.00 | -0.24 | 0.00 | 0.00 | 0.11 | -0.30 | 0.44 | 0.22 | 0.33 | -0.28 | 0.00 | 0.00 | 0.15 | 0.17 | -0.33 |
| 9 | 0.20 | -0.24 | 0.00 | 0.50 | 0.00 | -0.36 | 0.42 | -0.22 | 0.24 | 0.00 | 0.00 | -0.47 | 0.00 | 0.00 | 0.00 | 0.00 | 0.00 | 0.00 | 0.00 | 0.00 |
| 10 | 0.20 | -0.22 | 0.00 | 0.30 | 0.00 | 0.00 | 0.12 | 0.19 | 0.00 | -0.10 | 0.67 | 0.42 | 0.00 | -0.30 | 0.00 | 0.00 | 0.15 | 0.00 | 0.00 | 0.00 |
| 11 | 0.21 | -0.16 | 0.11 | 0.13 | 0.00 | 0.67 | 0.34 | 0.11 | -0.29 | -0.31 | -0.20 | -0.12 | 0.00 | 0.21 | 0.00 | -0.17 | 0.00 | 0.00 | -0.11 | 0.00 |
| 12 | 0.24 | 0.00 | 0.46 | 0.00 | -0.19 | 0.37 | 0.00 | 0.00 | 0.54 | 0.19 | 0.00 | 0.00 | 0.00 | -0.27 | 0.00 | 0.26 | -0.13 | 0.10 | 0.17 | -0.12 |
| 13 | 0.26 | 0.00 | 0.50 | 0.00 | -0.15 | -0.18 | -0.11 | -0.17 | 0.00 | 0.00 | 0.14 | 0.00 | -0.23 | 0.27 | -0.32 | -0.39 | 0.24 | -0.21 | -0.13 | 0.21 |
| 14 | 0.25 | 0.12 | 0.46 | 0.00 | -0.12 | -0.36 | 0.00 | 0.00 | -0.44 | -0.36 | 0.00 | 0.00 | 0.18 | 0.00 | 0.33 | 0.19 | -0.19 | 0.11 | 0.00 | -0.13 |
| 15 | 0.25 | 0.22 | 0.13 | 0.00 | 0.48 | -0.14 | 0.00 | 0.49 | -0.13 | 0.31 | -0.16 | -0.12 | 0.00 | -0.19 | -0.12 | 0.00 | 0.28 | 0.27 | -0.16 | 0.00 |
| 16 | 0.25 | 0.26 | 0.00 | 0.00 | 0.55 | 0.00 | 0.00 | 0.00 | 0.11 | 0.00 | 0.00 | 0.00 | 0.00 | 0.12 | 0.00 | 0.00 | -0.48 | -0.48 | 0.23 | 0.00 |
| 17 | 0.25 | 0.28 | -0.18 | 0.00 | 0.25 | 0.13 | -0.15 | -0.48 | 0.23 | -0.28 | 0.00 | 0.13 | 0.00 | 0.00 | 0.17 | 0.00 | 0.23 | 0.40 | -0.27 | 0.00 |
| 18 | 0.25 | 0.30 | -0.23 | 0.00 | -0.15 | 0.00 | 0.00 | -0.23 | -0.18 | 0.00 | 0.00 | -0.14 | 0.37 | -0.43 | -0.32 | -0.26 | 0.12 | 0.00 | 0.38 | 0.00 |
| 19 | 0.26 | 0.31 | -0.25 | 0.00 | -0.36 | 0.00 | 0.00 | 0.11 | 0.00 | 0.11 | 0.00 | 0.00 | 0.00 | 0.11 | -0.10 | 0.34 | 0.00 | -0.40 | -0.49 | -0.26 |
| 20 | 0.26 | 0.30 | -0.27 | 0.00 | -0.38 | 0.00 | 0.00 | 0.32 | 0.00 | 0.00 | 0.00 | 0.00 | -0.32 | 0.23 | 0.27 | -0.17 | -0.14 | 0.31 | 0.28 | 0.24 |

| NON ACTIVE | | | | | | | | | | | | | | | | | | | | |
|---|---|---|---|---|---|---|---|---|---|---|---|---|---|---|---|---|---|---|---|---|
| **bins** | **PC1** | **PC2** | **PC3** | **PC4** | **PC5** | **PC6** | **PC7** | **PC8** | **PC9** | **PC10** | **PC11** | **PC12** | **PC13** | **PC14** | **PC15** | **PC16** | **PC17** | **PC18** | **PC19** | **PC20** |
| 1 | | | | | | | | | | | -0.25 | -0.49 | -0.30 | 0.32 | | 0.19 | 0.65 | 0.18 | | |
| 2 | | | | | | | | | | | -0.33 | 0.57 | -0.23 | 0.20 | | 0.33 | | -0.24 | | 0.55 |
| 3 | | | | | | | | | | | -0.42 | 0.13 | | -0.25 | 0.19 | -0.75 | 0.28 | | | 0.24 |
| 4 | | | | | | | | | | | -0.27 | -0.40 | | 0.54 | 0.17 | -0.24 | -0.55 | -0.21 | 0.17 | 0.15 |
| 5 | | | | | | | | | | | -0.23 | 0.16 | | | -0.28 | | -0.19 | 0.70 | 0.54 | |
| 6 | | | | | | | | | | | -0.29 | 0.34 | 0.36 | 0.37 | 0.31 | | 0.22 | -0.12 | | -0.60 |
| 7 | | | | | | | | | | | -0.35 | -0.28 | 0.46 | -0.37 | 0.34 | 0.44 | -0.12 | 0.14 | -0.24 | 0.22 |
| 8 | | | | | | | | | | | -0.34 | -0.15 | 0.29 | -0.18 | -0.72 | | | -0.44 | | -0.11 |
| 9 | | | | | | | | | | | -0.33 | | -0.59 | -0.44 | 0.23 | 0.13 | -0.16 | -0.19 | 0.29 | -0.36 |
| 10 | | | | | | | | | | | -0.30 | | -0.25 | | -0.26 | | -0.28 | 0.30 | -0.72 | -0.26 |
| 11 | | -0.12 | 0.25 | | | -0.89 | -0.34 | | | | | | | | | | | | | |
| 12 | -0.17 | -0.59 | 0.69 | 0.11 | -0.22 | 0.29 | | | | | | | | | | | | | | |
| 13 | -0.30 | -0.61 | -0.33 | -0.38 | 0.51 | -0.12 | | | | | | | | | | | | | | |
| 14 | -0.34 | -0.24 | -0.53 | 0.19 | -0.60 | 0.13 | -0.36 | | | | | | | | | | | | | |
| 15 | -0.35 | | -0.12 | 0.45 | | -0.26 | 0.74 | | -0.16 | -0.14 | | | | | | | | | | |
| 16 | -0.36 | 0.15 | | 0.39 | 0.39 | 0.10 | -0.17 | 0.58 | 0.33 | 0.24 | | | | | | | | | | |
| 17 | -0.36 | 0.19 | 0.12 | 0.24 | 0.29 | 0.11 | -0.37 | -0.51 | -0.51 | | | | | | | | | | | |
| 18 | -0.36 | 0.21 | 0.11 | -0.20 | | | -0.11 | | 0.26 | -0.83 | | | | | | | | | | |
| 19 | -0.36 | 0.21 | 0.14 | -0.30 | -0.12 | | 0.17 | -0.44 | 0.53 | 0.45 | | | | | | | | | | |
| 20 | -0.36 | 0.24 | 0.11 | -0.50 | -0.25 | | | 0.46 | -0.49 | 0.17 | | | | | | | | | | |

**Table S2. Loadings for all Principle Components for active and non-active moths.** Bins from 1-20 refer to 1 s before to 1 s after the stimulus, stimulus started after bin 10. Absolute in- or decrease in between consecutive loadings reflect in- or decrease in flight strength. Missing values indicate values below an absolute value of 0.1. Colour-code reaches from dark blue (maximum value) to dark red (minimum value).

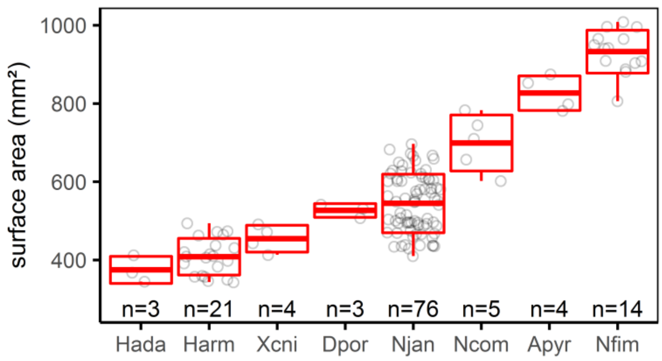


**Figure S1. Surface area of all moths with surface area measurements (N=130).** We assigned the mean surface area of each species to those individuals without surface area measurements. Boxplots show mean, standard deviation, and minimum/maximum value. Species abbreviations: Hada = *Heliothis adaucta*, Harm = *Helicoverpa armigera*, Xcni= *Xestia c-nigrum*, Dpor= *Deilephia porcellus*, Njan = *Noctua janthe*, Ncom= *Noctua comes*, Apyr= *Amphipyra pyramidae*, Nfim = *Noctua fimbriata*.

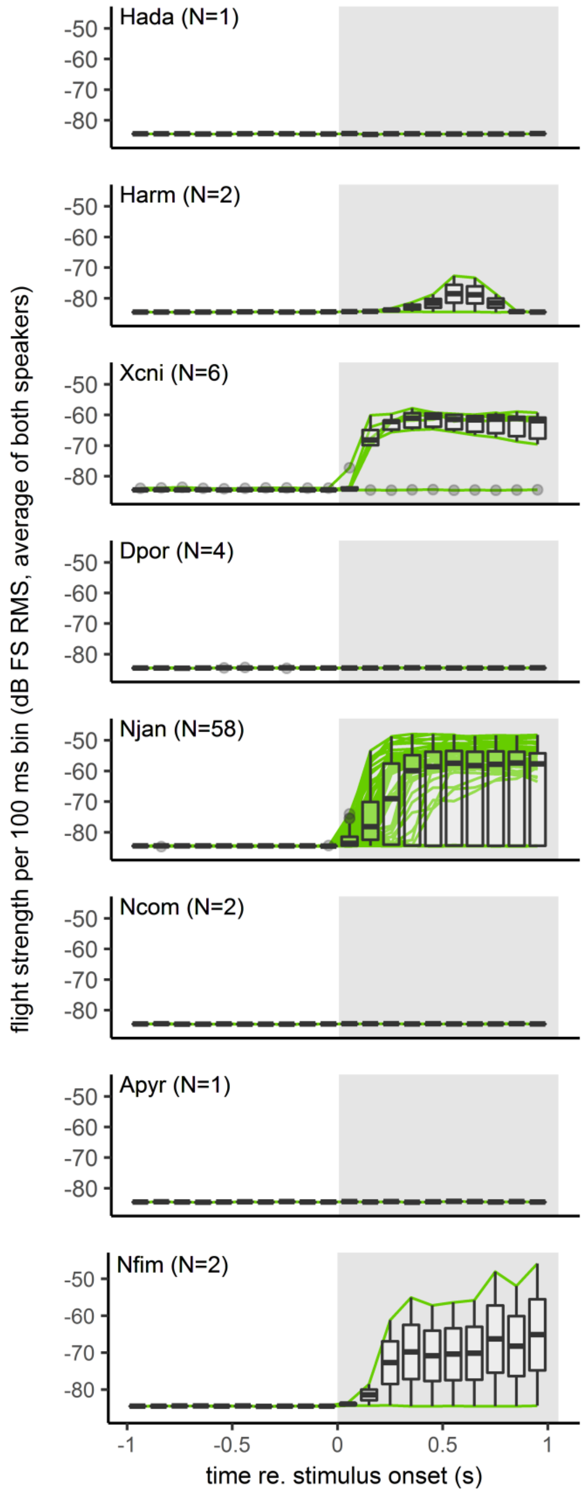


**Figure S2.** Flight strength per 100 ms bin for active moths from 1 s before to 1s after the stimulus onset for 8 moth species. Species and sample size are shown in the left upper corner of each panel. Boxplots show median, quartiles, whiskers (1.5 x inter-quartile range) and outliers beyond the interquartile range of the individual data (green lines). Stimulus presentation is indicated by the grey box. Species abbreviations: Hada = *Heliothis adaucta*, Harm = *Helicoverpa armigera*, Xcni= *Xestia c-nigrum*, Dpor= *Deilephia porcellus*, Njan = *Noctua janthe*, Ncom= *Noctua comes*, Apyr= *Amphipyra pyramidae*, Nfim = *Noctua fimbriata*.

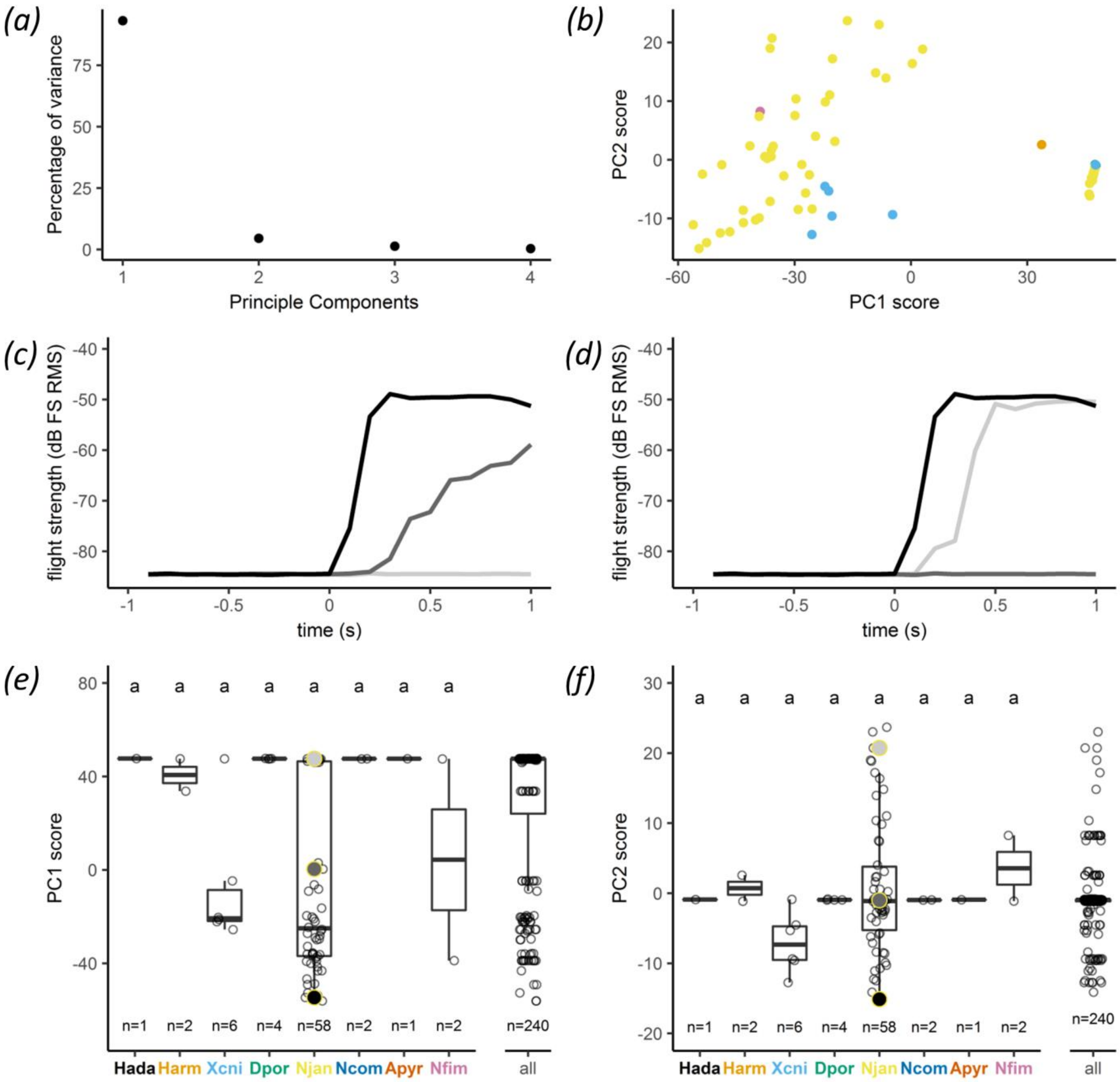


**Figure S3: Principle components of moths flying before stimulus onset ("active" moths).** (*a*) Percentage of variance explained by the first four principle components. (*b*) PC1 scores as a function of PC2 scores; for species colour code see panels e) and g). (*c*, *d*) Example flight behaviours for high (light grey), medium (dark grey) and low (black) values for PC1 and PC2. (*e*, *f*) PC1 and PC2 scores for each species and for the whole population ("all"), which was created by randomly sampling 30 values from each species. Species are ordered with increasing surface area from left to right. Letters above boxes indicate significant differences. Circled data points refer to examples shown above. Boxplots show median, quartiles, whiskers (1.5 x inter-quartile range) and outliers beyond the whiskers.
